# Automated microinjection for zebrafish xenograft models

**DOI:** 10.1101/2024.08.13.607596

**Authors:** Yi Ding, Kees-Jan van der Kolk, Wietske van der Ent, Michele Scotto di Mase, Saskia Kowald, Jenny Huizing, Jana M Vidal Teuton, Gunja Mishra, Maxime Kempers, Rusul Almter, Sandra Kunz, Laurine Munier, Carl Koschmann, Sebastian M. Waszak, Vincenzo Di Donato, Sylvia Dyballa, Peter Ten Dijke, Camila Vicencio Esguerra, Lasse D. Jensen, Jan de Sonneville

**Affiliations:** Life Science Methods BV, Leiden, the Netherlands; Centre for Molecular Medicine Norway (NCMM), University of Oslo, Oslo, Norway; ZeClinics SL, Barcelona, Spain; BioReperia AB, Linköping, Sweden; Oncode Institute, Dept. Cell & Chemical Biology, Leiden University Medical Center, Leiden, the Netherlands; Department of Pediatrics, University of Michigan, Ann Arbor, Michigan, USA; Division of Diagnostics and Specialist Medicine, Linköping University, Linköping, Sweden

**Author notes:** Shared first authors (Yi Ding, Kees-Jan van der Kolk).

## Abstract

Zebrafish xenograft models have been increasingly recognized for their ability to predict patient responses to cancer therapeutics, suggesting their potential as diagnostic tools in clinical settings. However, these models require the precise microinjection of cancer cell suspensions in many small and fragile zebrafish larvae. Manual injections are so challenging that, even after months of training, variability in experimental results persists among researchers. This limits the uptake and deployment of zebrafish xenograft models for clinical use and drug discovery. To address this challenge, we have designed, built, and validated an automated microinjection robot. Combined results of injections into the vasculature, perivitelline space, and hindbrain ventricle demonstrated an average injection success rate of approximately 60%, with a larvae survival rate exceeding 70%, comparable to manual injections using a traditional micromanipulator. Notably, the full automated mode was twice as fast as manual injections. This automation of the microinjection process significantly reduces the need for extensive personnel training while it enhances reproducibility, efficiency, and accuracy, paving the way for more extensive use of zebrafish xenograft models in drug discovery and patient diagnostics.

## Introduction

Over the past decades, the zebrafish (Danio rerio) has emerged as a powerful vertebrate model organism for studying various human diseases, including cancer (Choi et al., 2021; Liu & Leach, 2011). The introduction of human cells into zebrafish larvae, i.e., xenograft transplantation, is a widely employed technique to study tumor behavior in vivo, offering a dynamic and versatile model for cancer research (Chen et al., 2021; Gamble et al., 2021). Zebrafish larvae offer unique advantages over cell cultures and mammalian models in tumor research: their small size, high fecundity, transparency, tolerance for xenografts, the small number of cells required for implantation, and the rapid growth of xenografts. Zebrafish xenograft models have been used effectively to investigate processes such as tumor formation, migration, metastasis, intravasation, and extravasation (Fazio et al., 2020; Somasagara & Leung, 2022). Various cancer cell lines, including those from prostate cancer (Xu et al., 2018), breast cancer (Drabsch et al., 2013), urinary bladder cancer (Kowald et al., 2023), colorectal cancer (Fontana & Van Doan, 2024), glioblastoma (Pliakopanou et al., 2024), and pediatric brain tumors (Basheer et al., 2022) have been shown to form tumors, proliferate and/or extravasate in zebrafish. Additionally, the zebrafish patient-derived xenograft (PDX) model has gained traction as a promising model for predicting individual treatment responses and clinical outcomes in patients with colorectal (Costa et al., 2024), pancreatic (Barroso et al., 2021), epithelial ovarian cancer (Lindahl et al., 2024), non-small cell lung cancer (Ali et al., 2022) and rectal cancer (Costa et al., 2022).

Microinjection is a critical technique for introducing exogenous substances into zebrafish larvae for disease studies, such as cancer and infections. Zebrafish larvae present with multiple sites, which are well-tolerated injection locations including the duct of Cuvier (DoC), the perivitelline space (PVS), and the hindbrain ventricle. Systemic injection via the DoC allows introduction of the injected material directly into the bloodstream. This facilitates the in vivo evaluation of various stages of tumorigenesis, such as migration, intravasation, extravasation, and metastatic outgrowth at distant sites in cases where tumor cells are injected (Cabezas-Sáinz et al., 2020; Konantz et al., 2019; Mercatali et al., 2016; Pontes et al., 2017). Once in the bloodstream, different cancer cell subtypes, albeit inefficiently, survive, and intravasate in the tail region, with labeled cells being trackable and quantifiable (Kawakami et al., 2016). The PVS, located between the periderm and the yolk syncytial layer, is another commonly used injection site (Nicoli & Presta, 2007). The avascular nature of the PVS makes it ideal for studying newly formed blood vessels and further exploring migration, metastatic behavior, and tumor intravasation efficiency (Brown et al., 2017; Cabezas-Sáinz et al., 2020; Drabsch et al., 2017). The hindbrain ventricle is also an intriguing site. Injection of human melanoma cells and breast cancer cells into the hindbrain ventricle of zebrafish larvae has resulted in the formation of tumor masses and new vessels in the brain region (Gopal et al., 2023; Haldi et al., 2006). Additionally, the transplantation of glioblastoma cells into this site has shown increased tumor area, proliferation, migration, and responsiveness to chemotherapeutics (Rudzinska-Radecka et al., 2021; Wehmas et al., 2016).

Despite the invaluable insights derived from injections at these anatomical sites, the process of microinjection remains a daunting task for researchers that lack experience with these techniques. Conventionally, microinjection has been conducted across research laboratories worldwide using manual systems consisting of a stereo dissecting microscope, an air-pressure based microinjector, and a micromanipulator (Vagionitis Stavros & Czopka Tim, 2018; Q. Xu, 1999). This manual approach presents formidable challenges, mainly attributed to the tiny size and fragility of zebrafish larvae, the minimal needle tip size to fit cancer cells, and the complexities associated with targeting specific injection sites without causing overt damage to the larvae. Consequently, this manual process demands extensive training and practice, requires substantial time to perform the injections, and therefore has limited throughput capacity. Even after extensive training, variability in experimental results persists among researchers, arising from differences in skill levels, cognitive abilities, protocol interpretation, instrument usage, environmental conditions, psychological factors and individual biological and health conditions.

These technical challenges hinder the adoption of zebrafish xenotransplantation models for fast-turnover screening of compound efficacy for patient tumor treatment or drug discovery. Therefore, automatization of the injection process would accelerate the pace of tumor progression assessment and contribute to the timely assessment of therapeutic intervention. Some research groups have made efforts to innovate larvae preparation (Ellett F & Irimia D 2017) and the design of robotic microinjection systems (Chi et al., 2022; Qian et al., 2022; Zhang et al., 2021, Guo et al., 2024). However, such systems were not validated with xenograft assays, nor tested by life sciences researchers. While aqueous liquids are relatively simple to inject due to the low viscosity and high homogeneity of the liquids, injecting living cells represent an enormous challenge. Cells are large, requiring significantly larger diameters of the injection needles, which in turn may lead to more damage and reduced survival of the larvae. Furthermore, cells are likely to sediment, clump together, and may be more temperature- and/or time-sensitive compared to injection of non-viable biological material, causing higher but heterogeneous viscosity, fluidity, and therefore injection volume at constant pressure.

In this study, as part of the Eurostars consortium ROBO-FISH project, Life Science Methods (LSM) designed and developed a specialized robotic system for zebrafish larvae xenotransplantation procedures. The injection robot was subsequently tested and validated by injection of various substances and cancer cells at multiple laboratories of start-ups and institutions across Europe. These include Bioreperia in Sweden, the Centre for Molecular Medicine Norway (NCMM) in Norway, LSM and Leiden University Medical Center (LUMC) in the Netherlands, and ZeClinics in Spain. Results from multiple labs showed an average injection success rate of approximately 60% and a survival rate exceeding 70% for the three injection sites using the robot. These outcomes are highly comparable to manual injections using a traditional micromanipulator. Moreover, the fully automated mode of the robot was on average twice as fast as manual injections. Therefore, the robot accelerates the injection process, simplifies this challenging task, and reduces labor intensity, allowing higher throughput. This facilitates the uptake of zebrafish tumor xenograft models at new sites, potentially even at sites with minimal or no prior experience with zebrafish work, such as cancer clinics.

## Results

### Overview of the design of the injection robot for zebrafish larvae

To accommodate microinjections at various anatomical sites, we have designed a robot as depicted in **Figure 1A**. This system comprises several components to facilitate precise and adaptable injection processes. From an overhead perspective, we have incorporated a top-camera (acA1920-40uc, Basler AG, Germany) equipped with a high-resolution 10× telecentric lens with liquid focus (VS-TM10-55CO-LQL1, VST Europe B.V., the Netherlands), enabling detailed visualization of the injection procedures from above. To switch focus between the needle tip and the sample below, an Optotune liquid lens with an increased range was used (EL-16-40-TC-VIS-20D, Optotune Switzerland AG). In the middle section of the robot, the needle is affixed to a holder capable of rotation around the upper lens (**Supplementary Video 1**). This permits adjustment of the needle orientation to the target, a departure from the conventional approach of manipulating the sample or sample holder to match the needle orientation. The lower section of the system employs a camera (acA4024-29um, Basler AG, Germany) in conjunction with a 0.5× telecentric lens (VS-TCT05-65CO/S, VST Europe B.V., the Netherlands) to facilitate the detection of subsequent zebrafish larvae for injection. For illumination, a dome light, emitting broad rays of light, is employed to produce high numerical aperture (NA) lighting, resulting in a minimal depth of view as observed from the top-camera perspective, thereby enhancing surface focus (**Supplementary Video 2**). Additionally, a coaxial light source utilizes the input of the bottom lens to generate parallel, low-NA light, effectively increasing the depth of field (**Supplementary Video 3**) for the top camera. This feature proved invaluable, for instance, in the examination of intricate structures such as zebrafish blood flow. **Figure 1B** shows an image of the front side of the robot, presenting the fully integrated product. At the top, the black cylinder contains the camera, lenses, and a motor for rotating the needle at various angles. In the middle, two plates are located on the motorized stage: one 6-well plate for droplet calibration and an agarose gel plate for placing zebrafish larvae. At the bottom, there is a power button and a touch screen. The touch screen is used for the operation of the system and observation of the injection procedure in real-time.

**Figure 1.**
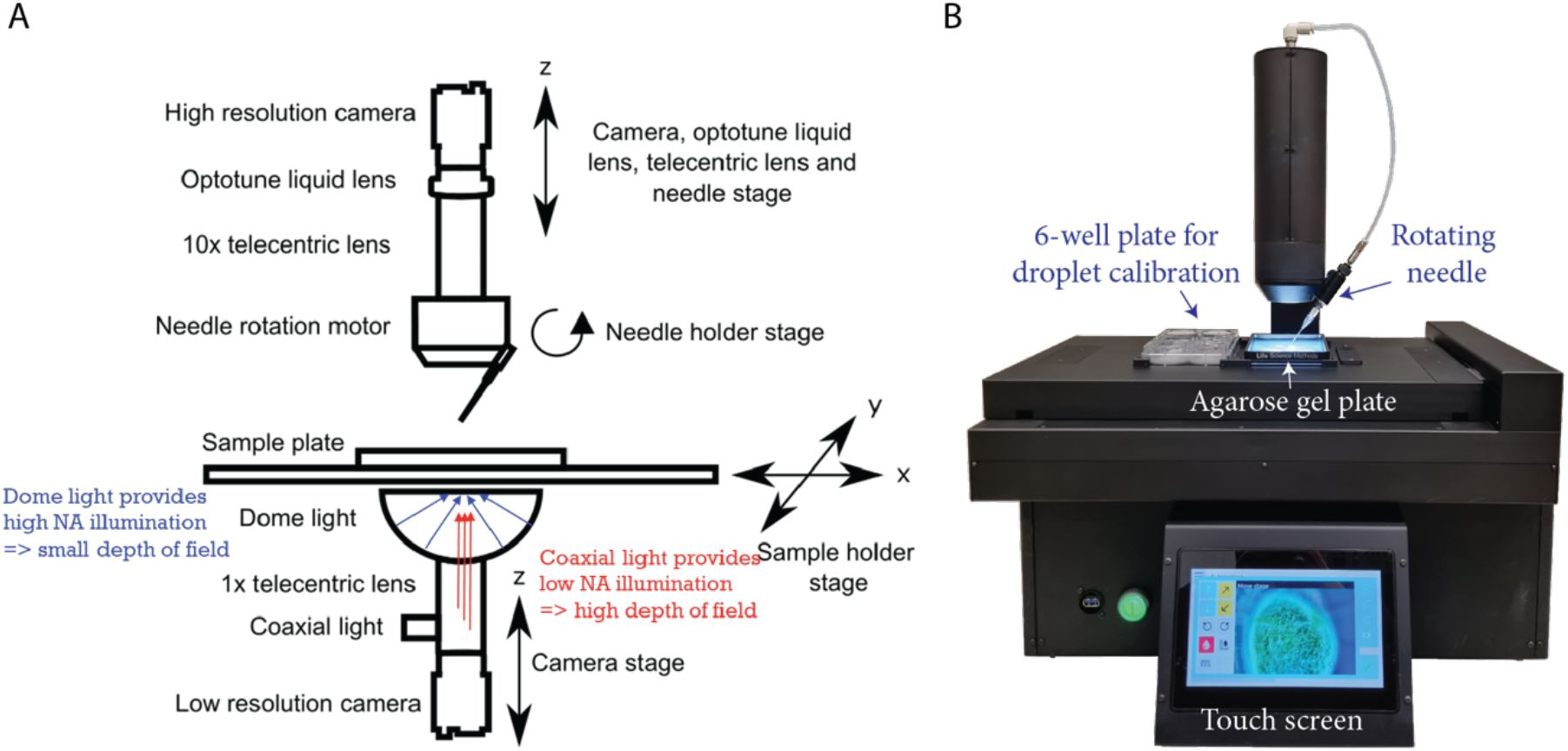
Design and visualization of the automated injection robot for zebrafish larvae. A. Schematic diagram illustrating the design and key components of the automated injection robot. B. Photograph showing the fully assembled automated injection robot in operation.

### Robotic injection procedure and puncture detection

The injection procedure for zebrafish larvae comprises five steps: injection settings, needle calibration, droplet calibration, plate selection, and injection. The procedure begins with configuring the injection settings, which include selecting different developmental stages, injection sites, injection locations with a specific needle starting point and angle as well as injection macros (**Figure 2A** and **Supplementary Video 4**). Next, needle calibration is performed to ensure the needle is at the correct height and validate the injection point of each rotation. This is achieved by adjustment of needle length and yaw (x, y) on the needle holder and focus (z) of the top camera using three screws on the robot (**Figure 2B** and **Supplementary Video 5**). The droplet size is then automatically measured in mineral oil within a 6-well plate. Injection pressure settings can be adjusted to achieve the desired volume (**Figure 2C** and **Supplementary Video 6**). In the plate selection interface, the stage can be moved into position to receive an agarose gel plate with around 20 anesthetized zebrafish larvae; these are randomly distributed (**Figure 2D** and **2E**). Finally, the injection process can commence (**Figure 2F**). To enable the robot to recognize the zebrafish larvae, thousands of images were collected and annotated (**Supplementary Figure S1**). This extensive dataset allows the larvae on the agarose plate to be accurately detected after the machine learning model is trained on the annotated zebrafish images (**Figure 2G**). The injection process begins with the robot scanning the agarose plate to locate a larva (**Figure 2H**). Once a larva is identified, the needle automatically navigates to the designated injection site. Using the liquid lens, the focus of the top camera is set to below the needle, and the head is lowered until the autofocus finds the larva. Subsequently, the focus is switched to the needle tip, and the approach continues until a touchdown is detected. Then, the injection macro is started if the automated mode is selected (**Figure 2H**). If not, the user can now take over control to guide an injection, which is the semi-automated mode. One strategy to detect puncture of the skin was to monitor the movement of labeled anchor points around the needle-tip (**Figure 2I** and **Supplementary Video 7**). Each anchor point is connected to a few pixels in the image; these are tracked, frame by frame. As the needle pushes the skin, the points move collectively in small increments. However, during puncture, a larger movement can be detected when the skin retracts more rapidly. Below, we show detailed injection results from three injection sites: duct of Cuvier, perivitelline space, and hindbrain ventricle.

**Figure 2.**
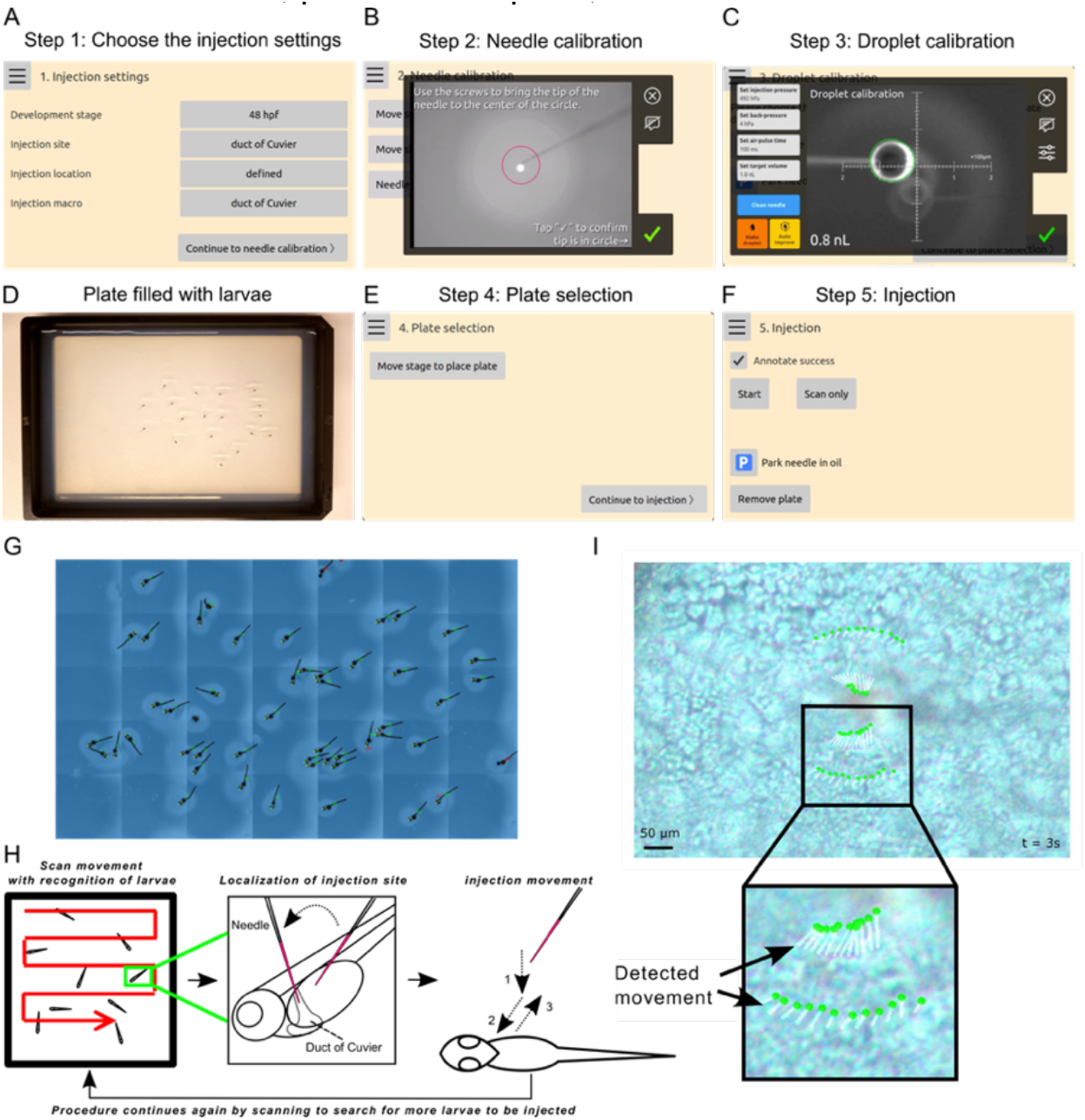
The injection procedure of the robotic injection system for zebrafish larvae. **A**.Various injection settings can be selected, including the developmental stage, injection site, injection location on schematic drawing, and injection macro. **B**.Calibration of the needle is performed in the x, y, and z directions in semi-automated mode. **C**. The volume of the injection droplet is automatically measured in a well filled with mineral oil. **D**. Prepare a 1.5% agarose gel plate and randomly place the anesthetized zebrafish larvae on the gel. **E**. Click “Move stage to place plate” and the robot will automatically move the stage to the correct position for placing the plate with the anesthetized larvae on the designated holder. **F**. In the injection interface, click the “Start” button to begin the injection process. The “Scan only” button enables scanning and image collection of zebrafish larvae for annotations. The “Park needle in oil” option allows the needle to rest in oil, preventing dryness and blockage. **G**. Zebrafish larvae are automatically detected by the robot after training the algorithm. **H**. The injection procedure involves scanning the plate to locate a larva, approaching the larva, and executing the injection macro if the automatic mode is selected. **I**. Skin puncture by the needle is detected through the movement of the marked anchor points.

### Injections into the duct of Cuvier of zebrafish larvae

The duct of Cuvier (DoC), also known as the common cardinal vein, is a major venous structure with extensive blood flow in zebrafish larvae. It is a broad conduit on the embryonic yolk sac that collects blood from the anterior and posterior cardinal veins and directs it to the heart (**Figure 3A**). This vein allows for the direct injection of substances into the bloodstream. In the injection settings, the needle tip can be positioned anywhere within the DoC area with flexibility in angle selection. **Figure 3B** is an example where the needle is placed at the center of the DoC, aligned against the direction of blood flow, and made visible by use of the coaxial illumination. Since a static image cannot capture flowing blood, a short video consisting of 10 frames was recorded, highlighting differences in red to make the blood flow clearly visible, see **Figure 3C**. The duct of Cuvier (DoC) region was annotated, and a machine-learning model was trained on thousands of annotated images of DoC (**Figure 3C**). Consequently, the robot can accurately recognize the shape of the DoC (**Figure 3C**). Various substances, including dyes (trypan blue, dextran, and phenol red), tumor-infiltrating lymphocytes (TILs), the human bladder cancer cell line UM-UC-3, microspheres, and the human breast cancer cell line MDA-MB-231, were injected into the DoC to validate the robot across four independent labs in Europe. In the automated mode, larvae are injected in a fully automatic manner. The injection site is positioned in the middle of the DoC, and the substance is clearly visible entering the bloodstream immediately (**Supplementary Video 8**). In the semi-automated mode, the robot recognizes and approaches the larvae automatically, and users have the flexibility to control the needle, adjust the focus, and perform the injection using the touchscreen. The injection site is chosen on the border of the yolk for larger substances like MDA-MB-231 cells (**Supplementary Video 9**). This is because the DoC layer is so thin that larger cells are more likely to be injected into the yolk. As shown in **Figure 3D** and the three-dimensional reconstructed video in **Supplementary Video 10**, the injected substances were well distributed in the bloodstream of zebrafish larvae. Furthermore, the common manual injection method using a micromanipulator was performed by skilled researchers for comparison with the robotic injections. Detailed data about the injection time, success rate, and survival rate for the injections into the DoC are presented in **Figure 3E**. The injection of dyes and TILs using both modes of the robot achieved success rates exceeding 70% at both Bioreperia and ZeClinics, while manual injections of dyes at Bioreperia had a success rate of 50%. At NCMM, the success rates for robotic and manual injections of microspheres were 54% and 47%, respectively. For cancer cell injections, the success rate of the robotic method ranged from 53% to 61%, slightly lower than the 70% success rate of the manual method. Overall, the average success rates for both modes of the robot (63% and 71%) were higher than that of the manual injector (56%). The injection time per larva using the automated mode (43 seconds) was approximately half that of manual injections (84 seconds). Consequently, the average number of larvae injected within a given period was twice as high with the automated injection compared to manual injection. The manual injection rate is highly variable and dependent on the experience of different researchers. In contrast, the robotic injection rate is much less dependent on the individual performing the procedure. Although the survival rate for robotic injections was slightly lower than that for manual injections, this discrepancy was primarily due to an outlier in the UM-UC-3 cell injection data.

**Figure 3.**
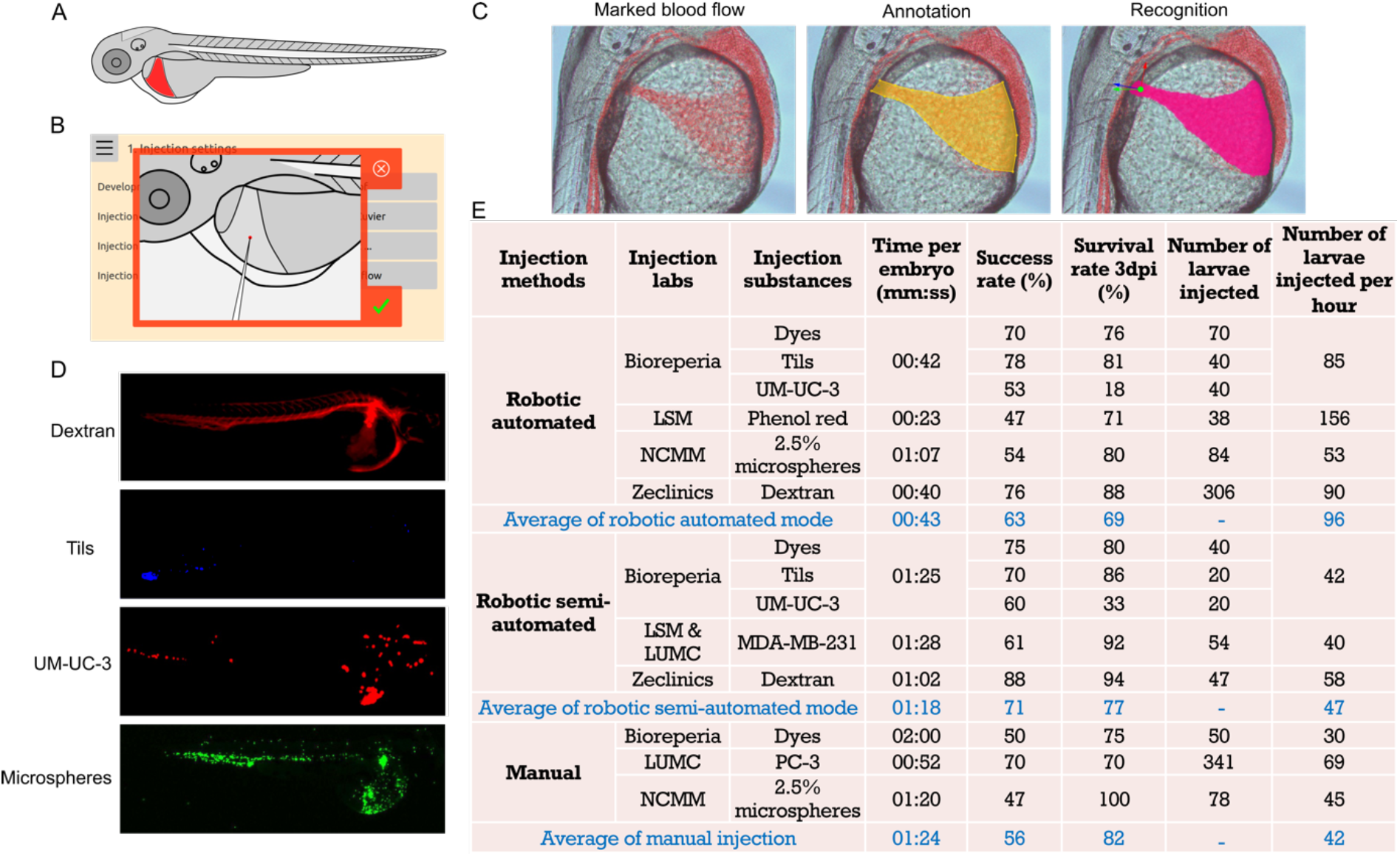
Injections into the duct of Cuvier (DoC) of zebrafish larvae. **A**. Schematic illustration of 2 days post-fertilization (dpf) zebrafish larvae, highlighting the targeted DoC area. **B**. Screenshot showing the needle angle and precise positioning of the needle tip for robotic DoC injections. **C**. Representative images demonstrating marked blood flow of 2 dpf larvae, annotation of the DoC, and recognition of the DoC by the robot after algorithm training on thousands of annotated images of DoC. **D**. Fluorescent images displaying the distribution of various substances within the bloodstream at 4 hours post-injection (hpi). TILs: tumor-infiltrating lymphocytes. **E**. Comprehensive data on the testing and validation of robotic DoC injections using different substances, with two robotic injection modes, conducted across multiple laboratories. Manual injections using micromanipulators were performed by different researchers for comparison to the robotic injections.

### Injections into the perivitelline space of zebrafish larvae

The perivitelline space (PVS) is an avascular area located between the periderm and the yolk syncytial layer (**Figure 4A**). For injection settings, it is recommended to position the needle tip at the upper border of the PVS with an angle of around 30 to 60 degrees (**Figure 4B**). To enable the robot to recognize the PVS structure, numerous images of the PVS area were collected, annotated, and used to train the algorithm (**Figure 4C**). With this trained algorithm, the robot can automatically navigate the needle to the predetermined starting point on the zebrafish larvae at the chosen angle. While MDA-MB-231 cells were predominantly used, other cancer cell types and clinical patient samples were also tested to validate these injections. Robotic automated and semi-automated injections of MDA-MB-231 cells into the PVS are shown in **Supplementary Videos 11** and **12**, respectively. Breast cancer cells were effectively delivered into the PVS at 4 hours post-injection (hpi) (**Figure 4D** and **Supplementary Video 13**). Cell migration was observed at 3 days post-injection (dpi). The quantification data of the three different injection methods for the PVS injections in multiple labs is presented in **Figure 4E**. For the automated mode, the success rate of MDA-MB-231 cell injections ranged from 39% to 62%. Using the semi-automated mode, the success rate for dyes, UM-UC-3, MDA-MB-231, and clinical samples was between 55% and 80%. With the manual injection method, the success rate for dyes, MDA-MB-231, and the human prostate cancer cell line PC-3 ranged from 70% to 75%. The success rate for HCC1806 cells was relatively low across all three injection methods performed at ZeClinics. Overall, the average injection success rate of the semi-automated mode (60%) was highly comparable to that of the manual methods (63%), while the automated mode had a relatively lower success rate (45%). However, the injection time using the automated mode was half that of the manual approach, allowing for more larvae to be injected within the same timeframe, which can compensate for the lower success rate. In terms of survival rate, the three injection approaches were highly similar, with the exception of a one-time discrepancy observed with MDA-MB-231 cells.

**Figure 4.**
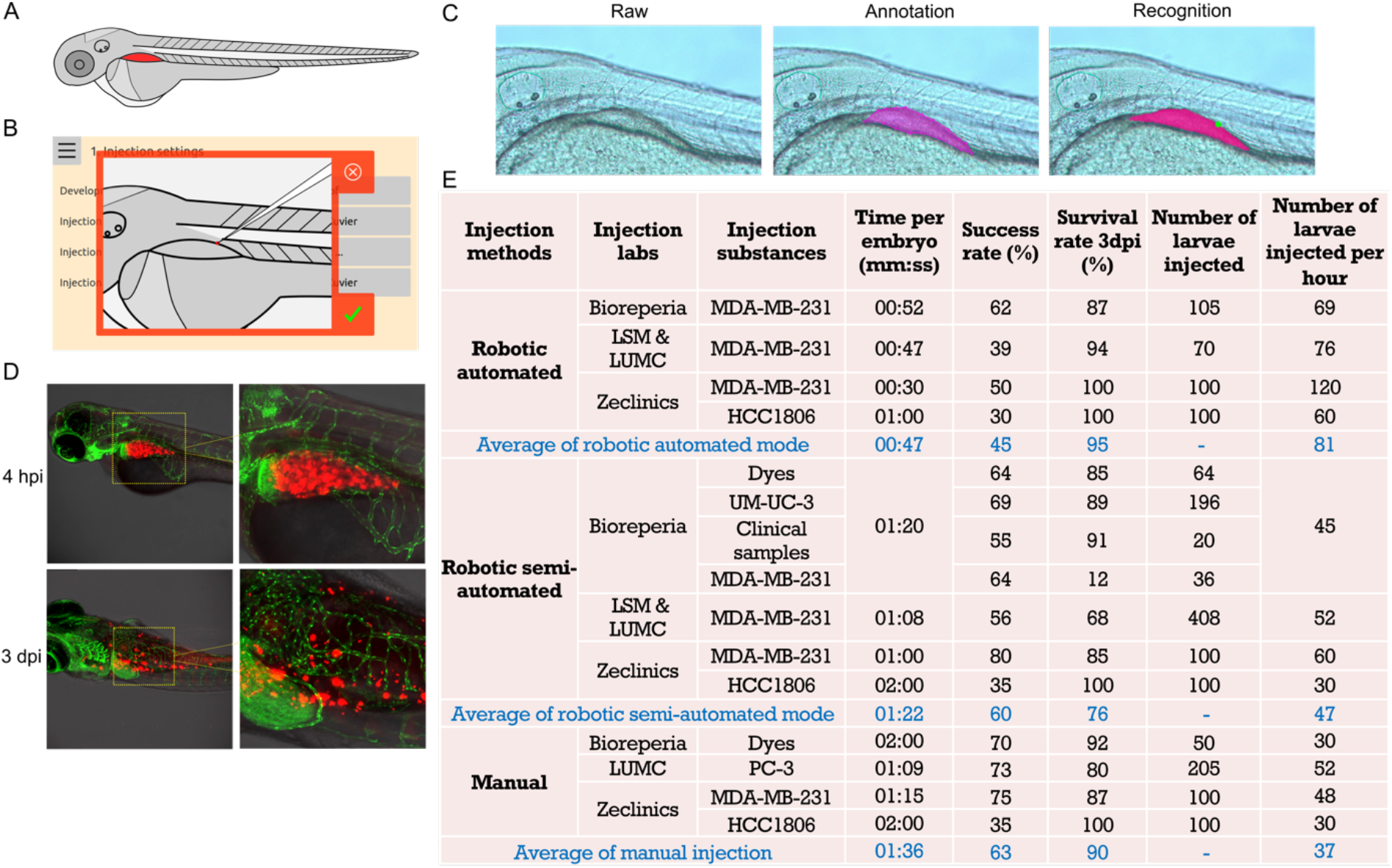
Injections into the perivitelline space (PVS) of zebrafish larvae. **A**. Schematic illustration of 2 days post-fertilization (dpf) zebrafish larvae, highlighting the targeted PVS area. **B**. Screenshot depicting the angle and precise positioning of the needle tip for robotic PVS injections. **C**. Representative images showing the raw images of the PVS acquired by the robot, annotation of the PVS, and recognition of the PVS by the robot after algorithm training on thousands of annotated PVS images. The green dot on the recognized image indicates the positioning of the needle tip. **D**. Fluorescent images displaying mCherry MDA-MB-231 cells in the PVS at 4 hours post-injection (hpi), and cell migration at 3 days post-injection (dpi). **E**. Comprehensive data on the testing and validation of robotic PVS injections using different cancer cell lines and clinical samples, with two robotic injection modes, conducted across multiple laboratories. Manual injections using a micromanipulator were performed as comparison to the robotic injections.

### Injections into the hindbrain ventricle of zebrafish larvae

The hindbrain in zebrafish larvae is anatomically located posterior to the midbrain and anterior to the spinal cord (**Figure 5A**). For injections into the hindbrain, the needle tip is typically positioned outside the hindbrain between the eye and otic vesicle, so that it punctures into the ventricle (**Figure 5B**). To enable the robot to detect the edge of the hindbrain, the borders of images of zebrafish larvae were annotated (**Figure 5C**). Additionally, the starting point and direction of the needle were annotated and trained to ensure automatic positioning at the predetermined site and angle (**Figure 5D**). Injection of the human GFP-labeled H3K27M-mutant diffuse midline glioma (DMG) cell line (SU-DIPG-XIII-P*) into the hindbrain ventricle resulted in cells being localized in the ventricle at 1-hour post-injection (hpi) and 4-day post-injection (dpi) (**Figure 5E**). The injection macro was developed after numerous manual injections and was optimized and validated by NCMM and LSM. In the automated macro mode, the robot will automatically place the needle, execute the macro, and perform the injection upon detecting the edge of the larvae (**Supplementary Video 14**). In the semi-automated macro mode, it requires clicking the macro in the manual interface, allowing flexibility for adjustments if needed (**Supplementary Video 15**). The quantification data using the two robotic modes was recorded at NCMM for injection of H3K27M-mutant DMG cells and at LSM for injection of the dye phenol red (**Figure 5F**). The success rates for dye and H3K27M-mutant DMG cell injections using the automated macro mode were similar (58% *vs* 57%). These rates were slightly lower than those achieved using the semi-automated macro mode (63%) and with manual injection (66%) for H3K27M-mutant DMG cells. However, the injection time of H3K27M-mutant DMG cells for the robotic automated macro mode was approximately half compared to the robotic semi-automated macro mode (120 *vs* 56 larvae/h) and manual injection (120 *vs* 57 larvae/h). All injection modes demonstrated high survival rates at 24 hpi (90-95%) (**Figure 5F**).

**Figure 5.**
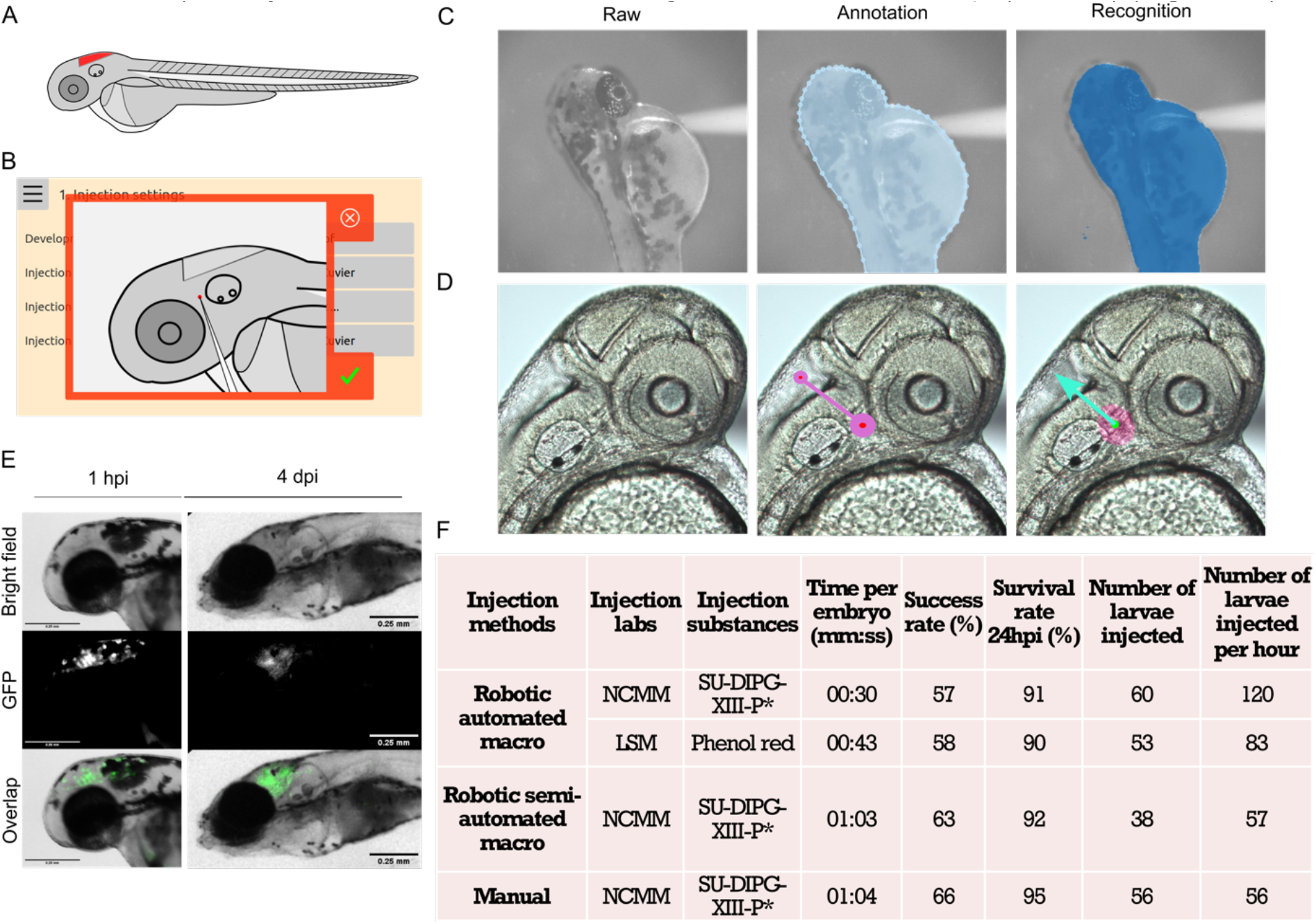
Injections into the hindbrain ventricle of zebrafish larvae. **A**. Schematic illustration of 2 days post-fertilization (dpf) zebrafish larvae, highlighting the targeted hindbrain ventricle. **B**. Screenshot showing the angle and precise positioning of the needle tip for robotic hindbrain injections. **C**. Representative images showing the raw images of 2 dpf larvae acquired by the robot, annotation of the larval edges, and recognition of larval shape by the robot after algorithm training on thousands of annotated images. **D**. The raw zoomed-in hindbrain images of 2 dpf larvae were acquired using the robot. These images were annotated with a line, where the large pink circle indicates the needle starting point, and the small pink circle indicates the needle direction. The needle starting point and direction then were recognized by the robot after algorithm training. **E**. Images displaying injected GFP-labelled H3K27M-mutant diffuse midline glioma cells (SU-DIPG-XIII-P*) into the hindbrain ventricle at 1-hour post-injection (hpi) and at 4 days post-injection (dpi). **F**. Comprehensive data on the testing and validation of robotic hindbrain ventricle injections using phenol red (LSM) and SU-DIPG-XIII-P* cells (NCMM) with two robotic injection modes. Manual injection of SU-DIPG-XIII-P* cells using a micromanipulator was performed as a comparison to the robotic injection modes.

## Discussion

In this study, we have developed and introduced a microinjection robot for zebrafish larvae. Utilizing recorded users’ best practices, combined with machine learning, macros were developed for fully automated injections into three locations: the bloodstream (through the duct of Cuvier), the perivitelline space, and hindbrain ventricle. The robot can be used in automated or semi-automated mode: The automated mode offers high speed, minimal human interference, and a relatively high success rate, while the semi-automated mode provides flexibility for more insight and development of new injection sites via a user-friendly touchscreen interface. The efficiency of the robot has been tested and validated with various substances by different operators in multiple independent laboratories across Europe. These broad tests help us to identify any lab-specific biases or limitations and ensure the system’s reliability, adaptability, and real-world applicability under diverse conditions. The success and survival rates of robotic injections were highly comparable to those of manual injections. Additionally, the automatic injection method offered a significantly faster speed, being twice as fast as manual injections. Manual microinjection typically requires highly skilled personnel, extensive training, and significant practice to achieve a high success rate. In contrast, operating the robotic system required only a few hours of training, significantly reducing the learning curve and making the technique more accessible.

Intravenous injection is the most common administration route in both mice and human patients. The major vein used for such injections in zebrafish is the duct of Cuvier (DoC). Injections into the DoC have been shown to effectively deliver the injected material throughout the circulatory system (Isogai et al., 2001). This method allows researchers to inject cancer cells into the DoC and observe tumor cell dissemination and metastasis in real-time, offering insights that are directly translatable to mammalian models and human conditions (Costa et al., 2020). However, achieving intravenous administration in zebrafish larvae presents significant technical challenges, including the need for clear visibility and automatic recognition of blood flow. The design of the coaxial light source, which generates parallel and low-numerical-aperture (NA) light, significantly increases the depth of field. This improvement makes the zebrafish blood flow visible, allowing the blood flow to be labeled and subsequently annotated and trained with an algorithm model. The trained algorithm greatly facilitates the recognition of the DoC and enables precise intravenous administration. Subcutaneous tumor implantation in mice allows for the development of localized tumors, which can be easily monitored and measured over time (Tomayko & Reynolds, 1989; Zhang et al., 2018). This method provides a controlled environment to study tumor growth, angiogenesis, and response to therapies (Ikeda et al., 2018). The perivitelline space (PVS) in zebrafish larvae is similar to the subcutaneous area in mice, making it a significant site for injections and localized tumor implantations. Microinjecting cancer cells into the zebrafish PVS is challenging due to its narrowness. In the automatic injection mode, the robotic system detects the needle’s skin puncture automatically. Additionally, the subsequent retraction of the needle in the macro ensures proper positioning within the PVS, leading to accurate cell delivery. The zebrafish brain, which is well-developed early in its development, shares the same major structural components as the mammalian brain, including the hindbrain (Guo, 2009; Kozol et al., 2016; Wilson et al., 2002). Hindbrain injections in zebrafish larvae are the most commonly used site for orthotopic implantation of brain tumor cells and for studying tumors that metastasize to the brain (Maricic et al., 2022). Since the hindbrain is located at the border of a larva, the edge detection design in the injection macro ensures precise needle placement within the hindbrain ventricle, allowing for accurate injection. H3K27M-mutant diffuse midline glioma (DMG) is a high-grade glioma typically diagnosed in children and young adults and preferentially located in the brainstem (pons, midbrain, medulla), thalamus, and spinal cord (Mackay et al., 2017). Surgical resection is limited due to anatomical location and standard treatment protocols (eg, chemotherapy) are ineffective, resulting in very poor clinical outcomes with 2-year overall survival rates of 10% (Hoffman et al., 2018). Novel drugs and combination therapies (Kline et al., 2022) are hence urgently needed for this patient population and our injection results of H3K27M-mutant DMG cells into the hindbrain ventricle of zebrafish larvae hold promise for future high-throughput drug screening efforts.

In its current status, the robotic system proved versatile, effectively handling multiple types of tumor cells, indicating its broad applicability in cancer research. The automation of xenotransplantation procedures not only enhanced precision but also significantly reduced the time required for these procedures, thereby increasing throughput, and facilitating large-scale studies. Post-injection analysis confirmed that tumor cells remained viable and proliferated within the zebrafish larvae, demonstrating successful xenotransplantation and the system’s suitability for subsequent research. Retrospective clinical trials have shown promising results, where patient tumor cells were tested in zebrafish xenograft models. These experiments have demonstrated a high correlation (up to 91%) between quantified tumor behavior in zebrafish and patients’ responses to standard clinical care for various types of cancers (Barroso et al., 2021; Costa et al., 2022, 2024). This high level of correlation indicates that xenograft assays offer a promising avenue for personalizing care, potentially reducing suffering and costs associated with multiple types of human cancers. Despite these advantages, the technique currently requires highly skilled personnel and years of practice, making it unsuitable to meet the demands for the number of patients, such as those with colorectal cancer (Morgan et al., 2023). Our previous work has demonstrated that automation can reduce variability and manual bias while increasing the yield compared to manual microinjections in zebrafish eggs (Cordero-Maldonado et al., 2019; Del Prado et al., 2024). The zebrafish larvae injection robot presented in this study has the potential to be implemented in diagnostic labs, such as pathology units or clinical genetics labs, within hospitals. This would enable the decentralization of functional precision medicine, making it accessible to a larger proportion of cancer patients. By integrating such automated systems into clinical workflows, we can enhance the scalability and efficiency of personalized cancer treatments, ultimately improving patient outcomes and expanding the reach of precision medicine.

Additionally, the robotic system offers high versatility in terms of automation across various injection sites, different types of injection applications, and injections in other hosts. Developments are underway for automating injections into alternative sites, such as the caudal vein, swim bladder, and pericardial space. Moreover, the robotic injection system is not limited to zebrafish xenograft models; it can be applied to various other applications, including infectious disease studies, compound screening, and toxicology assays. The system also shows potential for use in other species, such as mosquito egg injections for malaria research. Furthermore, continuous improvements and optimizations are being made to the injection robot to achieve higher success rates and faster speeds, paving the way for more challenging technical solutions in biomedical studies. The advancements in this automated system highlight its potential to transform experimental methodologies and improve the efficiency and scalability of preclinical research.

## Methods

### Zebrafish breeding and husbandry

Zebrafish were bred and managed in individual labs at Linköping University (supplier of zebrafish embryos to BioReperia), Leiden University, the Norwegian Centre for Molecular Medicine (NCMM), and ZeClinics. All breeding and husbandry practices adhered to local animal welfare regulations and followed standard protocols (http://zfin.org). For this study, we used the 2-day post-fertilization (dpf) wild type ABTL, the transparent Casper, and the Tg(Fli: GFP) Casper zebrafish lines, the latter of which has GFP-expressing vasculature.

### Injection substances

A variety of substances were used for injections, including phenol red (Sigma-Aldrich, #114529), trypan blue (Thermo Fisher, #15250061), and red fluorescent dextran (Thermos fisher scientific, # D1818), the human muscle-invasive urinary bladder cancer cell line UM-UC-3 (ATCC, #CRL-1749), human tumor-infiltrating lymphocytes (TILs, purchased under a special partnership agreement from 4C Biomed), microspheres (Sigma-Aldrich, #L4530), breast cancer cell lines mCherry-labeled MDA-MB-231 (ATCC, HTB-26)), GFP-labeled MDA-MB-231(gently provided by Dr. Fernanda Raquel da Silva Andrade), mCherry labeled human prostate cancer cell lines PC3 (a kind gift from Dr. Maréne Landström), and GFP-labeled HCC1806 (gently provided by Dr. Fernanda Raquel da Silva Andrade), GFP-labeled H3K27M-mutant diffuse midline glioma cells (SU-DIPG-XIII-P*) (Venkatesh et al., 2019), and clinical samples from bladder cancer patients. The clinical samples were collected at Vrinnevi Sjukhuset under a certain ethical permission (Kowald et al., 2023). The preparation of these substances was conducted as follows: UM-UC-3, TILs, and clinical samples were prepared at Bioreperia; GFP-labeled HCC1806 was cultured and prepared at ZeClinics; GFP-labeled SU-DIPG-XIII-P* cells were cultured and prepared at NCMM; MDA-MB-231 was cultured and prepared independently at Bioreperia, Leiden University Medical Center (LUMC), and ZeClinics. Human cells were frequently tested for the absence of mycoplasma, and cell lines were authenticated by short tandem repeat profiling. Cells were incubated under standard cell culture conditions at 37°C in a humidified incubator with 5% CO2. The cell culture medium was refreshed every two to three days, and cells were split at appropriate ratios as needed. Cells at 80% confluence were harvested using 0.5% trypsin-EDTA (Biowest, #MS0158100U) or StemPro Accutase (Gibco, #A11105-01) and subsequently washed three times with 1× PBS (VWR, #E403-500). The cells were filtered using a 40-70 μM cell strainer (Corning #352340) before resuspending in either culture medium or 2% polyvinylpyrrolidone 40 (PVP40, Sigma, #102420477) to achieve an approximate density of 2.5 × 108 cells/mL for injection. UM-UC-3 cells were fluorescently labeled using 1,1’-dioctadecyl-3,3,3’,3’-tetramethylindocarbocyanine perchlorate (DiI, ThermoFisher, #D3899) according to the manufacturer’s instructions. TILs were labeled with 25 μM CellTracker™ Blue CMAC dye (Thermo Fisher, #C2110). SU-DIPG-XIII-P* cells were cultured in DMEM/F-12 basal medium (Gibco, #11330032) supplemented with 1% B-27 Supplement (50X) Minus Vitamin A (Gibco, #12587-010), 1% Penicillin/Streptomycin (Gibco, #151400122), 0.1% Heparin (StemCell Technologies, #7980), 0.02% Epidermal Growth Factor (EGF, Shenandoah Biotechnology, #SHBT100-26), and 0.02% human recombinant Fibroblast Growth Factor (FGF, Shenandoah Biotechnology, #SHBT100-146). SU-DIPG-XIII-P* cells were kept in suspension and treated with StemPro Accutase Cell Dissociation Reagent (Gibco, #A11105-01). Approximately 3×106 SU-DIPG-XIII-P* cells/ml were pelleted at 300 x g for 5 minutes, and resuspended in 25 μl PBS, to achieve a final concentration of approximately 40,000 cells/μl for injection into hindbrain ventricles. Cryopreserved biopsies from bladder cancer patients were treated and prepared for injection following previously described methods (Kowald et al., 2023).

### Injection preparations

A 1.5% agarose gel was prepared by melting 1.5 g of agarose (Sphaero, #D00247) in 100 mL of egg water followed by pouring the solution into the square plate provided with the robot by LSM, ensuring that the surface of the plate is fully covered. The prepared injection substances were mixed and 5-8 μL were transferred to individual glass capillary needles using microloader tips. For injections of dyes or small-sized cells (e.g., TILs), needles with a tip diameter of 5-10 μm were used. For injections of larger cells (e.g., MDA-MB-231), needles with a tip diameter of 20 μm were used. Both manually pulled needles and commercial needles (Clunbury Scientific LLC, #B100-58-6, #B100-58-20) were utilized in this study. Approximately 2 minutes prior to implantation, zebrafish larvae were anesthetized in 40 μg/mL tricaine (Sigma, #E10521-50). Anesthetized zebrafish larvae were placed randomly on the agarose gel plate, ensuring no contact between larvae, and excess water was removed. Typically, about 20 larvae were added per plate for each injection batch. The plate with the anesthetized larvae was placed on the right plate holder of the stage in preparation for injection.

### Image annotation and deep learning algorithms

For the automatic detection of larvae and injection sites, deep learning networks were employed. Numerous images of ABTL and Casper zebrafish larvae were collected using the scanning function in the robot’s injection interface. These images, including those of the zebrafish larvae and specific injection sites, were annotated using an annotation tool developed by LSM. Using this tool, different regions of a larva were marked with a class (segmentation). In a postprocessing step, an overlap of classes was resolved by choosing a final class for every pixel in the image. For example, if a pixel was annotated as both “larva” and “eye”, then we chose “eye” as the final class as it provides more specific information.

The annotated images were then used to train a deep-learning network. For image segmentation tasks, we used a U-Net network topology (Ronneberger et al., 2015). We used as input an array of shape W×H×C (width × height × number of color channels), and as a result an array of shape W×H×N (width × height × number of classes). During the inference phase, i.e., after the network has been trained and is being used to drive decisions, each of the W×H pixels in the input image is assigned a predicted class. This is done by selecting the position of the highest value from a vector of the N computed values.

For detecting the moment of puncture of the skin we use a classification network. A classification network typically takes an entire image as input and produces a single class as output. We used the Inception v3 network (Szegedy et al., 2016) as the main vehicle. For each task, whether segmentation or classification, a portion of the annotated images served as training data, while the remaining images were used for validation during the training phase. Wet lab, real, experiments were performed to select the model with the best performance. If the performance was poor, more images were recorded and annotated, and the relevant networks were re-trained on a larger dataset. Automated image augmentation (Shorten & Khoshgoftaar, 2019) was used to artificially increase the number of training images.

For training we used a Shuttle desktop PC with an Intel(R) Core(TM) i9-9900, 64GB of memory, and 1TB SSD, combined with a NVIDIA GTX 1070 with 8GB memory. Since a U-Net can occupy a lot of computer memory, it was often necessary to divide the images into smaller images and combine the segmentations back into a larger segmentation. Most trainings took overnight for testing, and multiple days for deployment. Here, it is important to note that a U-Net tends to make mistakes near the edge of its input, since less information is available in a neighborhood near the edge than in the center of the image. Therefore, when combining multiple smaller segmentations (304 pixels in width and height), it turned out to be advantageous to overlap them by 50 pixels in both directions.

We found another improvement by observing that while the U-Net can make mistakes, it produces different results when its input is transformed in various ways. We chose 6 trivial transformations that are rotations by multiples of 90 degrees and reflections. During inference, for each of the transformations, the input image is transformed, and a segmentation is computed, which is then transformed back. Then, for each pixel in the resulting 6 segmentations, a consensus is computed based on the 6 computed classes for that pixel.

Deep learning methods are only the first step in the decision-making process of the robot. We will now outline briefly how segmentation and classification are used to inject into the duct of Cuvier of a zebrafish larva using the following steps:

1. The plate is scanned globally using the bottom camera; each image is segmented using a U-Net, where the eyes, yolk, body and swim bladder (or perivitelline space) of each larva are recognized.
2. The centroids of the eyes, yolk and swim bladder are computed.
3. A skewed (non-orthogonal) coordinate system is constructed from these points.
4. Using the user-defined coordinates (as expressed in this coordinate system) we move to the global area of the duct of Cuvier.
5. Using the top camera, we take multiple, more detailed images at the same location and we combine these images into an image that reveals the blood flow in that area. In this operation, we take the difference between two subsequent images, take the absolute value for every pixel, add these results, and normalize to obtain the final image showing the blood flow. This blood flow is then colored red and overlaid onto the last image of the set.
6. This image is then segmented using another U-Net, where the duct of Cuvier area is recognized.
7. Since the U-Net can produce mistakes, more than one area can be found. We choose the area which has the most pixels.
8. Using the previously found coordinate system and the segmentation, the needle is moved to the part of the duct of Cuvier as selected by the user.
9. The robot now adjusts the liquid lens to focus far below the needle, and the system moves the needle down towards the sample while recording images with the top camera.
10. Using a classical algorithm, the sharpness of the images is tracked. When the sharpness reaches a local maximum, the descent of the needle is stopped.
11. The liquid lens is now adjusted to focus just below the needle, the camera selects a smaller field of view, and the needle is moved down in smaller steps. When a sudden increase in sharpness is detected (i.e., a threshold value is reached), the system assumes that the needle is touching the surface.
12. Deep learning is used to detect puncturing with the needle, as follows:
13. If the stage moves in X and Y while the manipulator moves down, then the net effect can be achieved that the needle is moving along its length with respect to the larva.
14. While moving along the length of the needle, a stream of grayscale images is recorded with the top camera.
15. For each triplet of subsequent images, a new image is constructed having three channels.
16. A deep learning classification network that has been trained on these images is used to determine the moment when the needle punctures the skin. To train this network, we annotated the moment of puncture in a similar stream of images, and we used the image triplets as input to the training algorithm.
17. After the skin is punctured, the robot quickly moves the needle out slightly along its length to reduce the likelihood of damage, reduce pressure, and reduce possible obstructions near the tip of the needle, allowing a substance to be injected.
18. An air-pressure pulse is given, and the injection is performed.

### Automated injection mode

In the fully automated mode, the robot performs the injections autonomously. The robot scans the agarose gel plate from top to bottom to locate a larva. Upon detecting a larva, the robot approaches it and positions the needle at the predetermined injection sites. The robot then executes a macro tailored to each injection site (**Supplementary Videos 8, 11**, and **14**). After completing the injection, the robot resumes scanning for the next larva.

### Semi-automated injection mode

In the semi-automated mode, users have the flexibility to control and adjust the injection process. The robot scans the plate and navigates the needle to the predetermined injection site of a larva. A manual interface appears, allowing users to adjust needle rotation, focus, magnification, injection pressure, and back pressure and to move the needle (**Supplementary Videos 9** and **12**). Additionally, users can activate the injection macro by clicking the “m” button, which automates the injection while maintaining flexibility (**Supplementary Video 14**).

### Manual injections

Manual injections are conducted using a pneumatic picopump or femtojet 4x and a manipulator. The specific procedures for manual injections into the DoC and PVS have been previously described in detail (Li et al., 2022).

### Microscope imaging

Transfer the injected zebrafish larvae to petri dishes containing egg water. If cells were injected, the larvae were maintained in an incubator at 33 °C; otherwise, they were kept at 28.5 °C. In Bioreperia, LUMC, and NCMM, images of injected zebrafish larvae from all injection sites were acquired at 1 to 4 hours post-injection (hpi) using a stereo fluorescent microscope. Additionally, larvae from PVS and hindbrain injections were also imaged at 3 days post-injection (dpi) and 4 dpi, respectively. In ZeClinics, larvae from DoC and PVS injections were imaged at 1 to 4 hpi using the VAST Bioimager system (Union Biometrica, US), and the images were subsequently reconstructed into three-dimensional videos.

## Acknowledgment

We gratefully acknowledge the Eurostars program for supporting our research through the ROBO-FISH grant (Grant Number: E! 114899). S.M.W. was supported by a Pioneer Project grant from the Norwegian Cancer Society (#254836). We extend our sincere thanks to Dr. Maréne Landström and Dr. Fernanda Raquel da Silva Andrade for generously providing the cell lines used in this study. We also express our deep appreciation to the caretakers in the fish facilities for their dedicated care of the zebrafish.

## Author contributions

J.d.S. conceptualized and designed the robotic system. K.J.v.d.K. developed the control hardware, software, and machine learning algorithms for the robot. Y.D., J.H., M.K. and K.J.v.d.K. carried out the image annotations. Y.D., K.J.v.d.K., W.v.d.E., M.S.d.M., S.Ko., J.H., M.K., V.D.D., and J.d.S. developed, validated, and optimized the injection macros of the robotic system. Y.D., W.v.d.E., M.S.d.M., S.Ko., J.M.V.T., and L.M. performed the robotic injections into the DoC. Y.D., M.S.d.M., S.Ko., J.H., and R.A. performed the robotic injections into the PVS. S.M.W. conceived and designed the glioma study. C.K. provided materials and designed the glioma study. S.Ku. performed glioma cell culture and prepared cells for injection into the hindbrain ventricle. W.v.d.E. and Y.D. performed the robotic injections into the hindbrain ventricle. W.v.d.E., M.S.d.M., S.K., G.M., and L.M. performed the manual injections. J.d.S., L.D.J., C.V.E., and S.D. conceived the project. J.d.S., L.D.J., C.V.E., S.M.W., P.T.D. and V.D.D. supervised the research. Y.D. made figures and videos. Y.D. and J.d.S. wrote the first draft of the manuscript, with all authors contributing to the critical review and approval of the final version.

