## Supplementary figures and images for "Automated microinjection for zebrafish xenograft models"

### Supplementary Figure S1.pdf

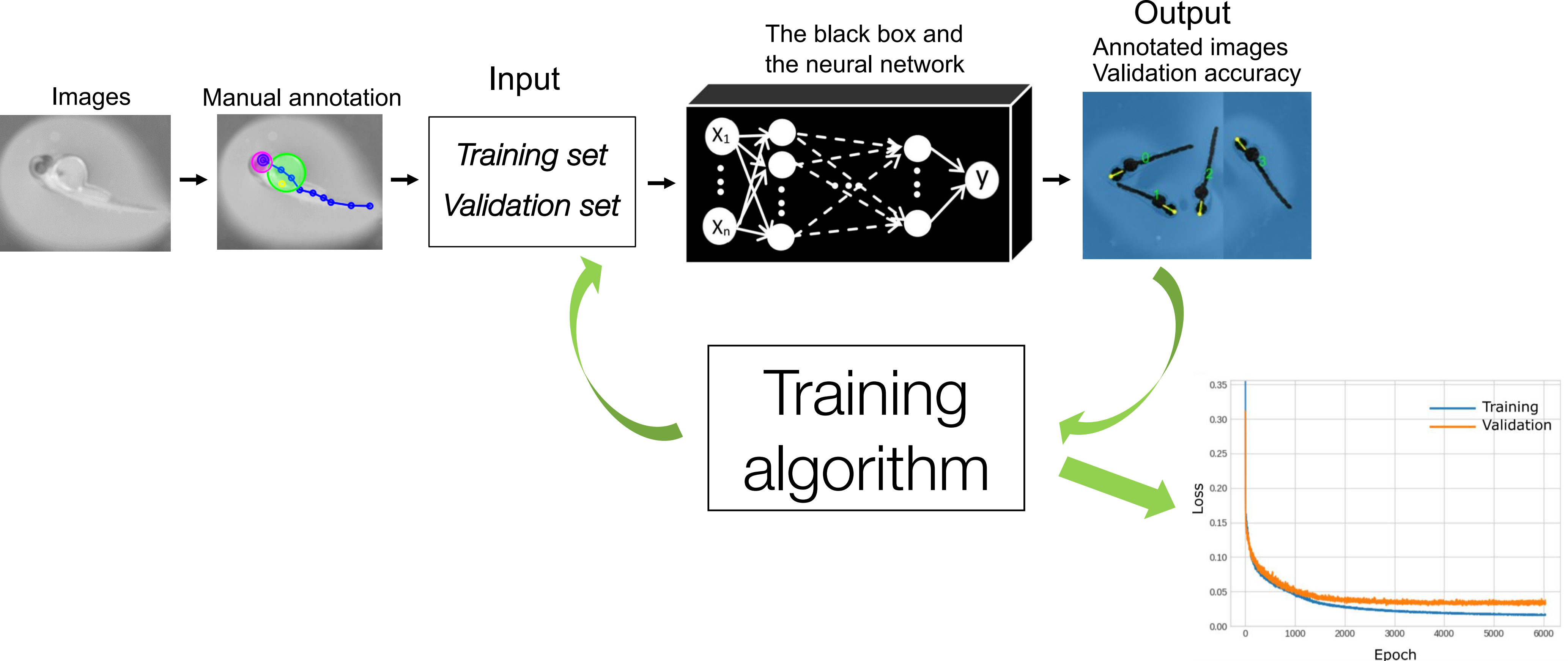
